# *In vivo* Perturb-Seq reveals neuronal and glial abnormalities associated with Autism risk genes

**DOI:** 10.1101/791525

**Authors:** Xin Jin, Sean K. Simmons, Amy X. Guo, Ashwin S. Shetty, Michelle Ko, Lan Nguyen, Elise Robinson, Paul Oyler, Nathan Curry, Giulio Deangeli, Simona Lodato, Joshua Z. Levin, Aviv Regev, Feng Zhang, Paola Arlotta

**Author notes:** Corresponding authors: Paola Arlotta, Feng Zhang, Aviv Regev, Xin Jin. These authors contributed equally (Aviv Regev, Feng Zhang, Paola Arlotta).

## Abstract

The thousands of disease risk genes and loci identified through human genetic studies far outstrip our current capacity to systematically study their functions. New experimental approaches are needed for functional investigations of large panels of genes in a biologically relevant context. Here, we developed a scalable genetic screen approach, *in vivo* Perturb-Seq, and applied this method to the functional evaluation of 35 autism spectrum disorder (ASD) *de novo* loss-of-function risk genes. Using CRISPR-Cas9, we introduced frameshift mutations in these risk genes in pools, within the developing brain *in utero*, and then performed single-cell RNA-Seq in the postnatal brain. We identified cell type-specific gene signatures from both neuronal and glial cell classes that are affected by genetic perturbations, and pointed at elements of both convergent and divergent cellular effects across this cohort of ASD risk genes. *In vivo* Perturb-Seq pioneers a systems genetics approach to investigate at scale how diverse mutations affect cell types and states in the biologically relevant context of the developing organism.

---

Human genetics has now uncovered strong associations between genetic variants in tens of thousands of loci and complex human diseases ranging from inflammatory bowel disease to psychiatric disorders (Jostins et al., 2012; Schizophrenia Working Group of the Psychiatric Genomics, 2014; de la Torre-Ubieta et al., 2016). In particular, analysis of trio-based whole-exome sequencing (WES) has implicated a large number of *de novo* variants in contributing to risk of several neurodevelopmental pathologies, including autism spectrum disorders (ASD) (Sanders et al., 2012; Satterstrom et al., 2018). Compared to common variants identified by Genome-Wide Association Studies, such *de novo* risk variants often have large effect sizes, are highly penetrant and are in the gene’s coding region, thus providing a crucial entry point for disease modeling and mechanistic studies. However, a major challenge remains for functional genetics: the identification of the point of action of these risk genes, each of which can, in principle, affect any of a massive number of different tissues, cell types, and molecular pathways. High-resolution and high-content phenotyping methods to identify tissue- and cell-type specific effects of genetic perturbations are needed, as generating and analyzing individual knockout animal models for long lists of risk genes as a first line of functional investigation is prohibitively time consuming and costly.

To address these challenges, we developed *in vivo* Perturb-Seq, a scalable, *in vivo* genetic screen, to investigate the function of large sets of genes at single-cell resolution in complex tissue *in vivo*. We applied *in vivo* Perturb-Seq *in utero* to study the effect of autism spectrum disorder (ASD) risk genes on mouse brain development. ASD comprises a broad collection of neurodevelopmental disorders with highly heterogeneous genetic contributions, including hundreds of highly penetrant *de novo* risk variant genes (Chen et al., 2015). Moreover, there is substantial diversity in the function of the gene products that risk genes encode, precluding a clear prediction about the underlying brain cell types, developmental processes, and molecular pathways affected during neurodevelopment (Mullins et al., 2016). By combining *in utero* genome editing in neural progenitors of the forebrain with postnatal single-cell RNA-Seq (scRNA-Seq), we studied how perturbing each of 35 ASD *de novo* variant genes affected brain development in a cell-type specific manner.

## *In vivo* Perturb-Seq to assess the function of ASD risk genes

We chose ASD candidate genes from a recently published WES study of 11,986 cases with 6,430 ASD probands, the largest published cohort in neural developmental disorder (NDD) and ASD genetics to date (Satterstrom et al., 2018) (**Table S1**). We initially prioritized 38 candidate genes (of which 35 were retained in the final analysis, **Supplemental Information**) that harbor a *de novo* variant specific to ASD patients within the broader class of neurodevelopmental disability (NDD) (**Figure S1A, Supplemental Information, Table S1**). These ASD risk genes are expressed in human brain tissue, as assessed by the Allen BrainSpan bulk RNA-Seq dataset (Miller et al., 2014); some are highly expressed at embryonic stages, and others highly expressed from early postnatal to adult stages (**Figure S1B**). Based on mouse cortical scRNA-Seq data, their orthologs are expressed in diverse cell types (**Figure S2**) (E18.5 data from the 10x Genomics public dataset, see **Supplemental Information**; P7 data from this work). Thus, these ASD genes could in principle act in many different cell types and temporal frames, emphasizing the importance of using scalable methods to test gene function across a range of cell types and developmental events.

For *in vivo* Perturb-Seq, we used Cas9-mediated genome editing (Adamson et al., 2016; Dixit et al., 2016; Jaitin et al., 2016) in a pooled approach to introduce mutations in each of the ASD risk genes within progenitor cells of the developing forebrain *in utero*, followed by scRNA-Seq at P7 to read out both a barcode of the perturbation and the perturbed cell’s transcriptomic profile (**Figure 1A**). Specifically, we used a transgenic mouse line that constitutively expresses Cas9 (Platt et al., 2014), and delivered pools of gRNAs by lentiviral infection into the lateral ventricles of the developing embryo *in utero*. Each lentiviral vector contained two gRNAs targeting the 5’-end coding exons of one ASD gene and a blue fluorescent protein (BFP) reporter with a barcode corresponding to the perturbation identity (Adamson et al., 2016; Dixit et al., 2016; Jaitin et al., 2016). To minimize vector recombination, we packaged each lentivirus independently and then pooled them at equal titers. We injected a pool of lentiviruses with equal gRNA representation into the ventricular zone at E12.5 (**Figure 1A**). In this approach, lentiviral infection will label neural progenitors lining the ventricle, and since the virus integrates into the genome, their progeny will likewise be labeled by BFP and carry a perturbation barcode corresponding to the targeted ASD gene. The lentiviral administration allows a sparse labeling of less than 0.1% of cells in the cortex (**Figure S3A-C)** On P7, we micro-dissected and dissociated cortical and striatal tissue, FACS-enriched the perturbed cells by selecting for BFP expression, and used massively parallel scRNA-Seq to obtain each cell’s expression profile along with its perturbation barcode. The cell survival rate after FACS was 78%, and we confirmed a 40-70% frameshift insertion/deletion for each gRNA target among the infected cells (**Figure S3D-E**).

**Figure 1.**
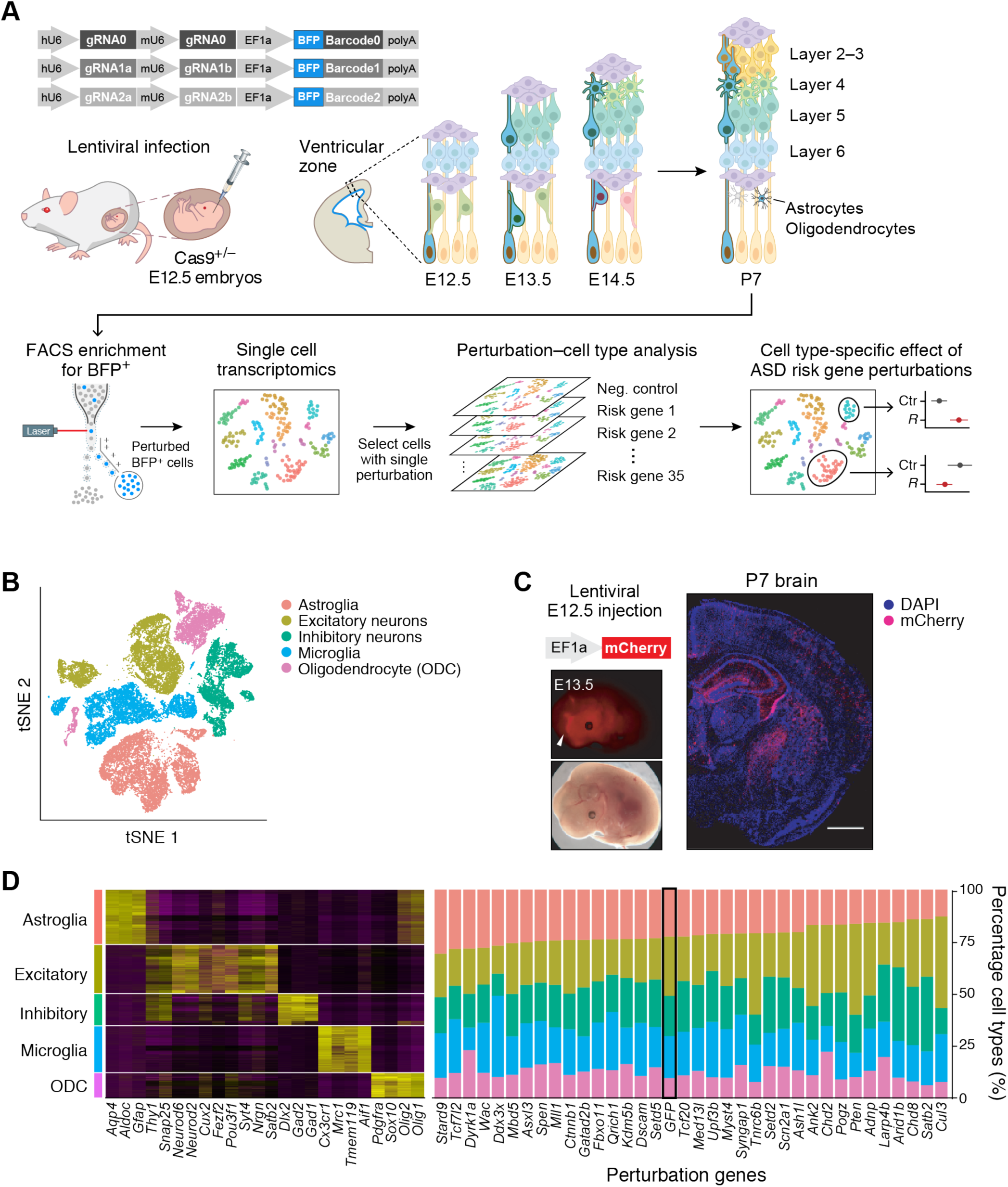
*In vivo* Perturb-Seq to investigate functions of a panel of ASD risk genes harboring *de novo* variants. (A) Schematics of the *in vivo* Perturb-Seq platform, which introduces mutations in individual genes *in utero* lentivirally at E12.5, followed by transcriptomic profiling of the cellular progeny of these perturbed cells at P7 via scRNA-Seq. (B) TSNE of five major cell types identified in the Perturb-Seq cells. (C) *In vivo* Perturb-Seq lentiviral vector with an mCherry expression starts within 24h, and can sparsely infect brain cells across many brain regions. Scale bar is 1000μm. (D) Cell type analysis of *in vivo* Perturb-Seq of ASD *de novo* risk genes. Canonical marker genes were used to identify major cell clusters (left), and cell type percentage representation in each perturbation group (right). Negative control (GFP) is highlighted by a black rectangle.

## *In vivo* Perturb-Seq targets diverse cell types without affecting overall cell composition

Targeting of the gRNA library to the lateral ventricle of E12.5 embryos results in infection of neural progenitors of the cortex, striatum and hippocampus. This allowed us to examine the effects of each perturbation across a wide range of progeny cell types (i.e. projection neurons, interneurons, astroglia, oligodendroglia, etc.) from distinct brain regions. In agreement immunohistochemical analysis and scRNA-Seq showed that the Perturb-Seq vector was expressed across a variety of neuronal and glial cell types (**Figure 1B, Figure S3A-B**).

We performed the experiment with 18 different cohorts of pregnant mice, for a total of 163 embryos, each subjected to the entire pool of perturbations. This multiplexed experimental design allowed us to test the cell-autonomous effect of all perturbations against a negative control construct targeting the endogenous GFP in the Rosa26 locus, a construct that was included in the same pool thus minimizing batch-dependent variation (**Figure S3F**). After quality control, we retained for further analysis a total of 46,770 cells from the neocortex across 17 high-quality experimental batches, and 7,118 striatal cells from 6 experimental batches. We grouped the cells into major subsets using Louvain clustering (Blondel et al., 2008) and annotated them by known marker gene expression (Zeisel et al., 2018) (**Figure 1D**). We then focused on five broad cell populations for downstream analysis: cortical projection neurons, cortical inhibitory neurons, astrocytes, oligodendrocytes, and microglia (thus excluding vascular, endothelial, hippocampal and striatal cells (**Figure S3F**)). After filtering some remaining low-quality cells in these groups (**Supplemental Information**), we retained 35,857 high quality cells (median of 2,436 detected genes per cell overall, and 4,084 genes in the projection neuron cluster (**Figure S3G**)). We subclustered each of these five major cell types separately and annotated biologically meaningful subclusters (**Figure S6**).

Based on the perturbation barcodes from the lentiviral constructs, 92% (33,231 cells) of the cells in these five major cell types had at least one perturbation read assigned to them (**Supplemental Information**), and 50% had a single gene perturbation identity (**Figure S4A-C**, 18,044 cells), reflecting the low multiplicity of infection (**Figure S4D**). We assigned 18,044 cells to a single perturbation at a median of 338 cells per perturbation. After excluding genetic perturbations with <70 perturbed cells detected, we retained 35 ASD risk gene perturbations in the final analysis. BFP from the lentiviral vector was robustly detected as one of the most highly expressed genes in all cells retained for analysis (**Figure S4E**), and the BFP detection rate in each cell type was correlated to the average number of genes detected (**Figure S4F**).

Relative to the negative control (which targeted the *GFP* gene), ASD risk gene perturbations had a very modest effect on the composition of the five main cell types. Only loss of *Dyrk1a* had a significant effect on cell type composition, increasing the proportion of oligodendrocytes and reducing microglia [(FDR-corrected *P*<0.05 using Poisson regression (Haber et al., 2017)] (**Figure 1D, Figure S5**).

## ASD gene perturbations affect gene programs and cell states within and across sub-populations of cells

To assess whether molecular changes and alterations in cell state were caused by ASD genetic perturbations, we next defined modules of genes within each of the five cell types that co-varied as a group across the cells within each cluster (**Figure 2A**). Such modules may reflect common biological processes (*e.g.*, cell cycle, differentiation, cell identity) whose activity either varies naturally across the cells within a given subset (Bielecki et al., 2018; Smillie et al., 2019), and/or is affected by the introduced perturbation. As previous work has shown (Adamson et al., 2016; Dixit et al., 2016; Jaitin et al., 2016; Duan et al., 2019), focusing on gene modules instead of individual genes provides more power to detect biologically meaningful perturbation effects using fewer cells than would be required for single gene-level analysis. To recover these gene expression modules, we applied two algorithms: Weighted Gene Correlation Network Analysis (WGCNA) and structural topic modeling (STM) (**Supplemental Information, Figure S6-7, Table S2-3**) (Roberts et al.; Langfelder and Horvath, 2008). As the modules selected by WGCNA were highly correlated with one or more topics (the STM analogue of modules) (**Figure S7**), we focused on 14 WGCNA modules extracted from the five major cell types.

**Figure 2.**
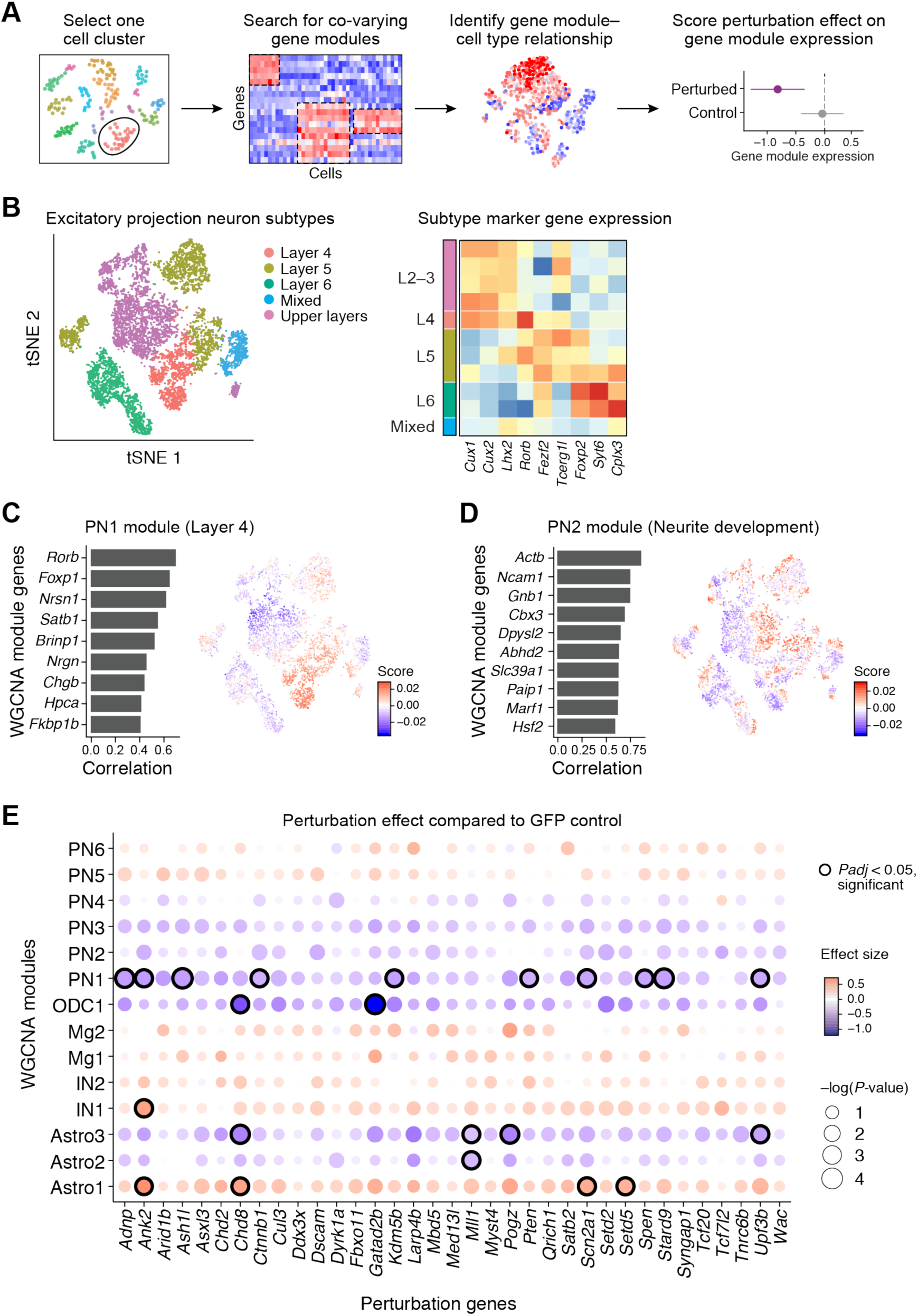
*In vivo* Perturb-Seq reveals cell-type specific effects of ASD risk gene perturbations. (A) Schematic illustration of the Perturb-Seq analysis pipeline, using co-varying gene module analysis in each cell cluster to estimate perturbation-associated effects on gene modules using linear modeling. (B) Subtypes of projection neurons (left), identified by expression of key marker genes (right). (C-D) Examples of projection neuron co-varying gene modules associated with Layer 4 projection neuron subtype identity and neurite development, respectively. (E) ASD risk gene perturbation effects in different WGCNA gene modules compared to GFP controls. Dot color corresponds to effect size, dot size corresponds to log(P-values). Module gene lists are in Table S2. *P-value* was calculated using a permutation-based approach; *Padj* was calculated using Benjamini & Hochberg FDR correction.

Within each cell category, some modules were specific to one subcluster within a cell type, whereas others ranged across cells in multiple subclusters, reflecting association with a specific subtype or a cell state, respectively. For example, within the projection neuron cluster, some of the modules are specific to subclusters matching a subtype of projection neurons, Layer 4 projection neurons (module PN1) (**Figure 2B**,**C**). Conversely, module PN2, associated with genes involved in neurite development and varied across cells in multiple subclusters, regardless of subtypes (**Figure 2D**).

We next tested whether perturbation of individual ASD risk genes was associated with changes of expression in each module. For each of the 14 WGCNA gene modules, we fit a joint linear regression model to estimate the effect size of each perturbation on that module. This allowed us to measure how module gene expression in each perturbation group deviated from the GFP control group (**Figure 2E**). We estimated the significance of these deviations by permuting the perturbation labels across cells within each experimental batch separately and comparing the resulting effect size to that in the unpermuted data. To ensure that no single perturbation or batch had a dominant effect on the linear model, we down-sampled cells in each cell category such that no perturbation had more than two times the median number of cells over all perturbations (**Supplemental Information**).

Perturbations in 15 ASD genes (*Adnp, Ank2, Ash1l, Chd8, Ctnnb1, Gatad2b, Kdm5b, Mll1, Pogz, Pten, Scn2a1, Setd5, Spen, Stard9*, and *Upf3b)* had significant effects across six modules (**Figure 2E**, circles, compared to the GFP control, FDR corrected *P*<0.05, **Table S4**): the projection neuron Layer 4 module (PN1), all three astroglia modules (Astro1, Astro2, and Astro3), the oligodendrocyte progenitor module 1 (ODC1), and the interneuron *Ndnf+* module (IN1). Perturbation of three additional ASD risk genes, *Ctnnb1, Dscam*, and *Setd2*, resulted in nearly significant (FDR corrected *P*<0.09) decreases in the projection neuron neurite development module (PN2) (**Figure 2D-E**). This is consistent with previous work showing that *Dscam* regulates presynaptic assembly and arborization size in *Drosophila* sensory neurons (Kim et al., 2013), and that loss of function of *Ctnnb1* signaling impairs synaptic vesicle diffusion, decreases vesicle number, and decreases dendritic arborization (Bamji et al., 2003; Yu and Malenka, 2003; Gao et al., 2007).

We further used the non-parametric van der Waerden test to ask which modules had a significant amount of their variation across the relevant cell types explained by the ASD perturbations overall. The oligodendrocyte progenitor module (ODC1) was a significant hit (FDR corrected *P*<0.05), while two other modules, Astro2 and Astro3 (corresponding to astroglia progenitors and astrocyte activation, respectively), were nominally significant (*P*<0.05, FDR corrected *P*>0.05) (**Figure S5C**). In order to determine whether changes in module expression observed in neocortical cells may also occur in cells of other brain regions, we performed *in vivo* Perturb-seq in the striatum and analyzed a total of 7,118 cells (5,933 of which are in glia clusters) from 6 independent batches. We find that the direction of effect of most perturbations largely agrees with those in the cortical data (see Supplemental text and **fig S10, Supplemental Information**), indicating that at least some of the ASD gene perturbation effects appear to generalize in cells of more than one brain region

Collectively, the data indicate that a selected group of perturbations was able to affect specific gene expression modules with cell-type specificity and point to convergent effects on such modules by a subset of perturbations.

## Single perturbation of *Ank2* confirms effect on interneuron gene expression module

In our multiplex *in vivo* Perturb-seq results, *Ank2* perturbation leads to an increase in the interneuron *Ndnf*+ module (IN1) (FDR corrected *P<*0.05, **Figure S8**). *Ank2* encodes an ankyrin protein, which interacts with ion channels and can stabilize GABAergic synapses (Tseng et al., 2015). To validate our finding, we tested this result in a simplex setting, by performing a single perturbation targeting either *Ank2* or *GFP* (control), followed by scRNA-Seq of neocortical cells at P7, collecting 2,943 and 1,716 high quality cells, respectively.

The individual simplex perturbation confirmed the results from the multiplexed *in vivo* Perturb-Seq screen. First, *Ank2*-perturbed cells were present across all cell types and their overall proportions were not significantly changed (**Figure S8B**). The IN1 module was strongly correlated with a subcluster of inhibitory neurons, which expressed *Ndnf* (**Figure S8C,D**), and within those cells *Ank2* loss-of-function perturbation led to upregulation of the module (FDR corrected *P<*0.05, **Figure S8E**), confirming the *in vivo* Perturb-Seq result.

## The ASD risk genes *Chd8* and *Gatad2b* alter gene programs in oligodendrocyte progenitors

*Chd8* and *Gatad2b* perturbations significantly decreased the expression of the ODC1 module in the oligodendrocyte cluster (**Figure 3A-D**, FDR corrected *P*<0.05). The ODC1 module is expressed highly in cycling cells and oligodendrocyte precursor cells (OPC), and lowly in newly formed oligodendrocytes (nODC) and myelinating oligodendrocytes (mODC), suggesting a link to oligodendrocyte maturation (**Figure 3A**).

**Figure 3.**
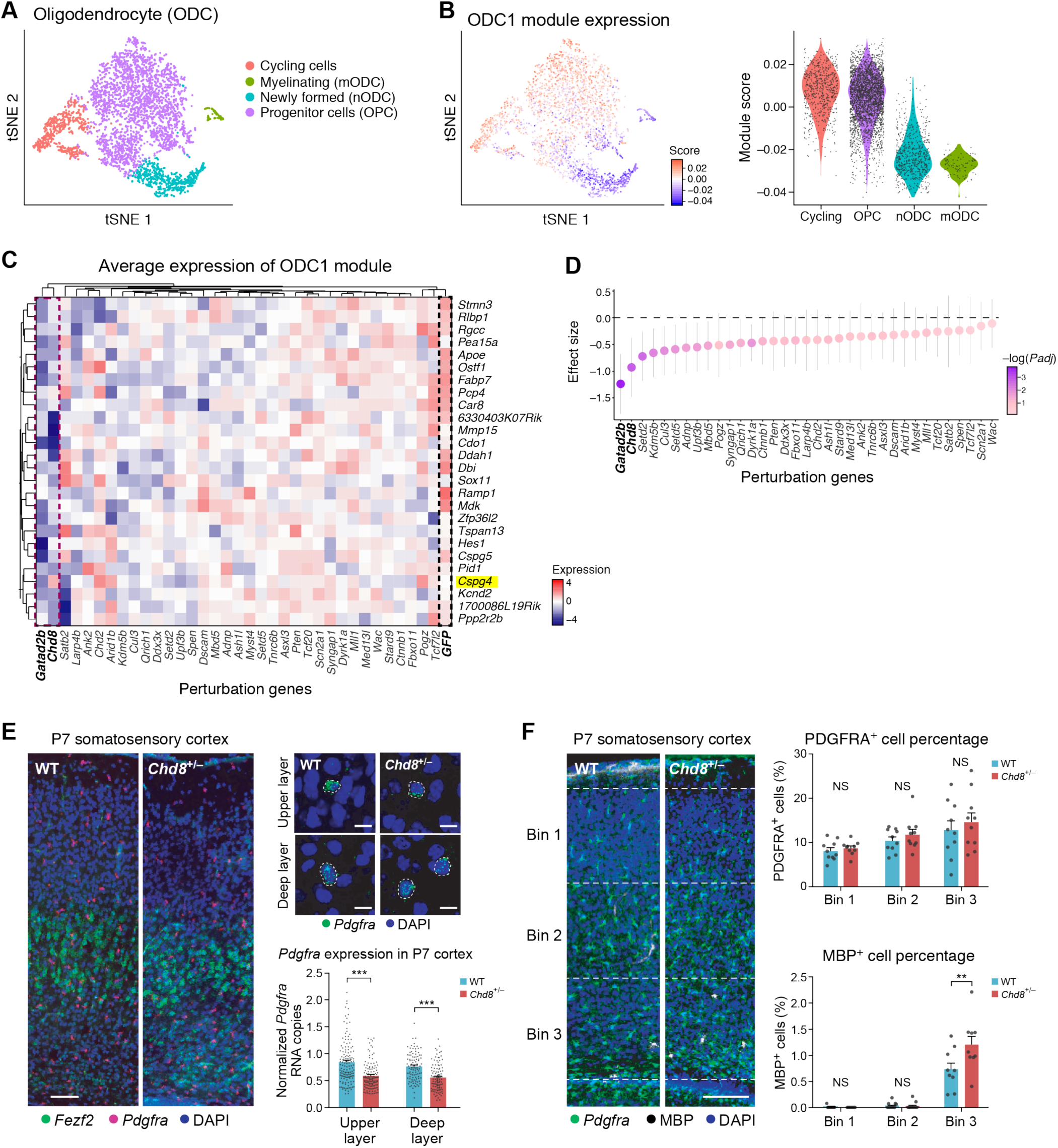
Perturbation effect in oligodendrocytes and validation in *Chd8*^*+/-*^ mouse models. (A) TSNE of oligodendrocyte subtypes from the Perturb-Seq data. (B) The ODC1 gene module expression score in each cell (left) and in each subcluster (right). (C) Average expression of genes in the ODC1 gene module (by row) in each perturbation group (by column), scaled by row. (D) Effect size of each perturbation on the ODC1 gene module compared to the control group. Error bars represent 95% confidence intervals. (E) *In situ* hybridization for *Pdgfra*, a marker of oligodendrocyte precursor cells (OPC), in the somatosensory cortex of P7 *Chd8*^*+/-*^ and wild-type littermates. Right: quantification of *Pdgfra* expression in P7 cortex of *Chd8*^*+/-*^ and wild-type littermates. Each dot represents the gene expression measurement from one cell; error bars represent standard error of the mean (n=2-3 animals per genotype). Scale bar on the left top panel is 1000μm, left bottom panel is 200μm and right panel is 50μm. (F) Immunohistochemistry for PDGFRA and MBP, markers for immature OPC and mature oligodendrocytes, and cell counts in the somatosensory cortex of P7 *Chd8*^*+/-*^ animals and wildtype littermates. Scale bar is 250μm.

We further investigated and validated this result by examining oligodendrocyte development in a *Chd8* germline heterozygous mutant model (as homozygous mutation is embryonic lethal (Nishiyama et al., 2004)), using several orthogonal methods. First, we used *in situ* hybridization for two canonical OPC marker genes, *Pdgfra* and *Cspg4*, one of which (*Cspg4*) is in the ODC1 module. Both were downregulated in P7 *Chd8*^*+/-*^cortex (**Figure 3E, Figure S9A-C**), consistent with our *in vivo* Perturb-Seq results. Immunohistochemistry against PDGFRA did not show significant differences in OPC cell numbers between the WT and *Chd8*^*+/-*^ littermates at P7 and P12, also consistent with *in vivo* Perturb-Seq; however, cells positive for the myelinating protein MBP were increased in numbers and displayed elevated MBP protein levels in the *Chd8*^*+/-*^ mutant in both P7 and P12 (FDR corrected *P*<0.05, nonparametric ANOVA test) (**Figure 3F, Figure S9D-G**). These results also agree with previous findings that *Chd8* loss of function is connected to abnormal OPC development (Marie et al., 2018). Collectively these data indicate that *in vivo* Perturb-Seq can identify cell-type specific molecular changes that agree with single gene perturbations.

## Discussion

*In vivo* Perturb-Seq can serve as a scalable tool for systems genetic studies of large gene panels to reveal their function at single-cell resolution in complex tissues. In this work, we demonstrated the application of *in vivo* Perturb-Seq to ASD risk genes in the developing brain; more generally, this method can be applied across diverse diseases and tissues.

ASD affects brain function profoundly, but its cellular and molecular substrates are not yet defined. The large number of highly penetrant *de novo* risk genes implicated through human genetic studies offer an entry point to identify the cell types, developmental events, and mechanisms underlying ASD origin. However, this requires scalable methods to define the function of genetic hits, with cell-type specificity. Here, we observed different cell types and processes affected by distinct ASD risk genes as well as distinct molecular pathways which are differentially affected across cell types. In addition to effects in neurons, oligodendrocyte development was affected by certain perturbations. Oligodendrocytes modulate and consolidate neural circuit refinement, and abnormal maturation of oligodendrocytes may be linked to long-lasting changes in neural wiring and brain function (Bercury and Macklin, 2015). One of the risk genes, *Chd8*, encodes a protein that binds directly to β-catenin and negatively regulates the Wnt signaling pathway, which plays a crucial role in progenitor proliferation and differentiation in the forebrain (Sakamoto et al., 2000; Durak et al., 2016; Katayama et al., 2016; Platt et al., 2017). Our results showed that *Chd8* modulates gene programs for oligodendrocyte differentiation and maturation, consistent with previously reported ChIP-Seq results showing that CHD8 interacts directly with OPC maturation genes in the neonatal stage (Marie et al., 2018; Zhao et al., 2018). Other cell types may be altered at different developmental stages, through cell-autonomous (intrinsic) or non-cell autonomous (extrinsic) mechanisms, which should be investigated further in the future.

Although we focused on the neocortex in this study, *in vivo* Perturb-Seq can be applied to study gene functions systematically across other tissues, to reveal tissue-specific as well as broadly-distributed gene functions, and uncover both the impact of individual disease-associated genes and the overall set of processes that they affect. Our findings underscore the importance of using single-cell profiles as a rich, comprehensive and interpretable phenotypic readout. With advances in other single cell profiling approaches (e.g., single-cell ATAC-Seq (Rubin et al., 2019)), single-cell multi-omics (Bian et al., 2018), and spatial genomics (Wang et al., 2018; Rodriques et al., 2019), we expect *in vivo* Perturb-Seq to be coupled in the near future with diverse readouts to better define the function of disease-risk associated variants from molecular mechanisms to non-cell autonomous effects in tissues. Spatial transcriptomics in particular should be well suited for use with *in vivo* Perturb-Seq, and should help uncover non-cell autonomous effects. *In vivo* Perturb-Seq can enable discoveries of pathways and cell types affected in heterogenous genetic pathologies, directing downstream studies and informing the development of refined models for genetic disorders and mechanistic studies as we move from genetic variants to function.

## Acknowledgement

We thank C. Dulac and M. Meselson for critical reading of our manuscript; Leslie Gaffney, Anna Hupalowska, Rhiannon Macrae, Juliana Brown, as well as members of the Levin lab, Regev lab, Zhang lab, and Arlotta lab for technical and intellectual support. This work is supported by the NARSAD Young Investigator Award and Harvard William F. Milton Grant (to X.J.); Stanley Center for Psychiatric Research at the Broad Institute (to J.Z.L.); The Klarman Cell Observatory, HHMI and an NHGRI Center for Cell Circuits CEGS grant (to A.R.); NIH grants (1R01-HG009761, 1R01-MH110049, and 1DP1-HL141201), HHMI, New York Stem Cell and Mathers Foundations, the Poitras Center for Affective Disorders Research at MIT, the Hock E. Tan and K. Lisa Yang Center for Autism Research at MIT, and J. and P. Poitras (to F.Z.); Stanley Center for Psychiatric Research at the Broad Institute, NIH grants (U01MH115727, R01MH096066, P50MH094271 Conte Center) (to P.A). F.Z. is a New York Stem Cell Foundation–Robertson Investigator. AR is a co-founder of and equity holder in Celsius Therapeutics and an SAB member of ThermoFisher Scientific, Syros Pharmaceuticlas and Neogene Therapeutics. F.Z. is a co-founder of Editas Medicine, Beam Therapeutics, Pairwise Plants, Arbor Biotechnologies, and Sherlock Biosciences. X.J., S.S., A.R., F.Z. and P.A. are co-inventors on *in vivo* Perturb-Seq and CRISPR inventions filed by the Broad Institute relating to the work in this manuscript. Data generated for this study are available through the Gene Expression Omnibus (GEO, accession #tbd) as well as the Broad single cell portal (#tbd). The analysis pipeline is deposited on KCO GitHub repository (https://github.com/klarman-cell-observatory/ivPerturbSeq). All other data are available in the manuscript or the supplementary materials.

## Supplemental Information

### Tables

Table S1. ASD risk gene list and their effect in ASD/NDD patient cohort.

Table S2. WGCNA gene module gene lists.

Table S3. Structural topic modeling fitted model.

Table S4. Effect size estimate of each ASD risk gene perturbation and nonparametric ANOVA analysis in the cortex in WGCNA modules and STM topics.

Table S5. Effect size estimate of each ASD risk gene perturbation and nonparametric ANOVA analysis in the striatum in WGCNA modules.

Table S6. gRNA design for the ASD risk gene perturbations.

Table S7. Parameters used in Seurat for cell type clustering.

### Figures S1-S10

**Figure S1.**
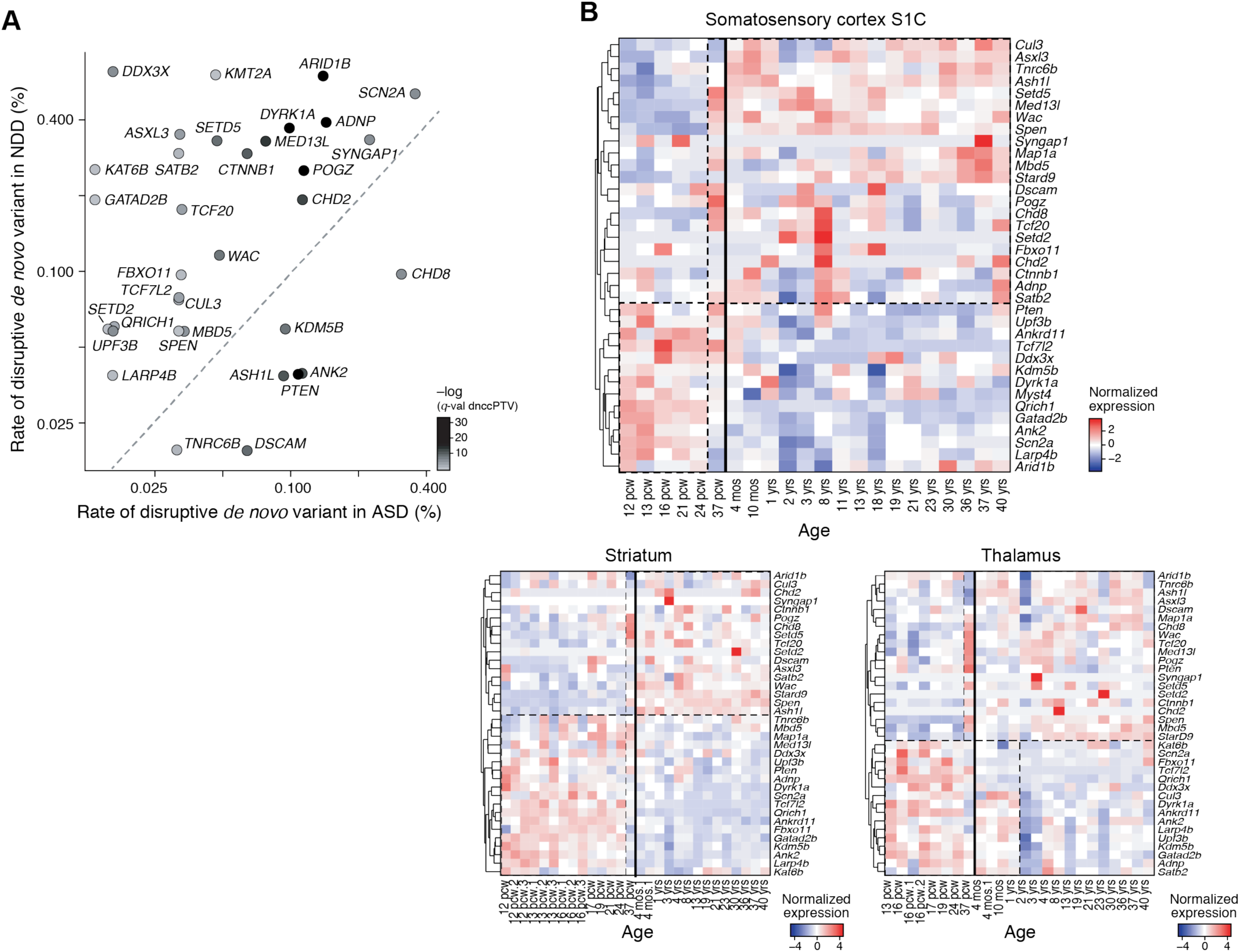
(A) The frequency of *de novo* loss-of-function variants in ascertained Autism Spectrum Disorders (ASD) and ascertained neurodevelopmental delay (NDD) cases for the 35 risk-associated genes included the Perturb-Seq analysis. *Q* value was calculated based on the de novo and case control (dncc) data. This data comes from Satterstrom *et al*. (B) Gene expression of a panel of selected ASD *de novo* risk genes in human somatosensory cortex (S1C), striatum, and thalamus across the Allen Brain Atlas BrainSpan postmortem samples from various ages. Dendrogram indicates hierarchical clustering by the rows.

**Figure S2.**
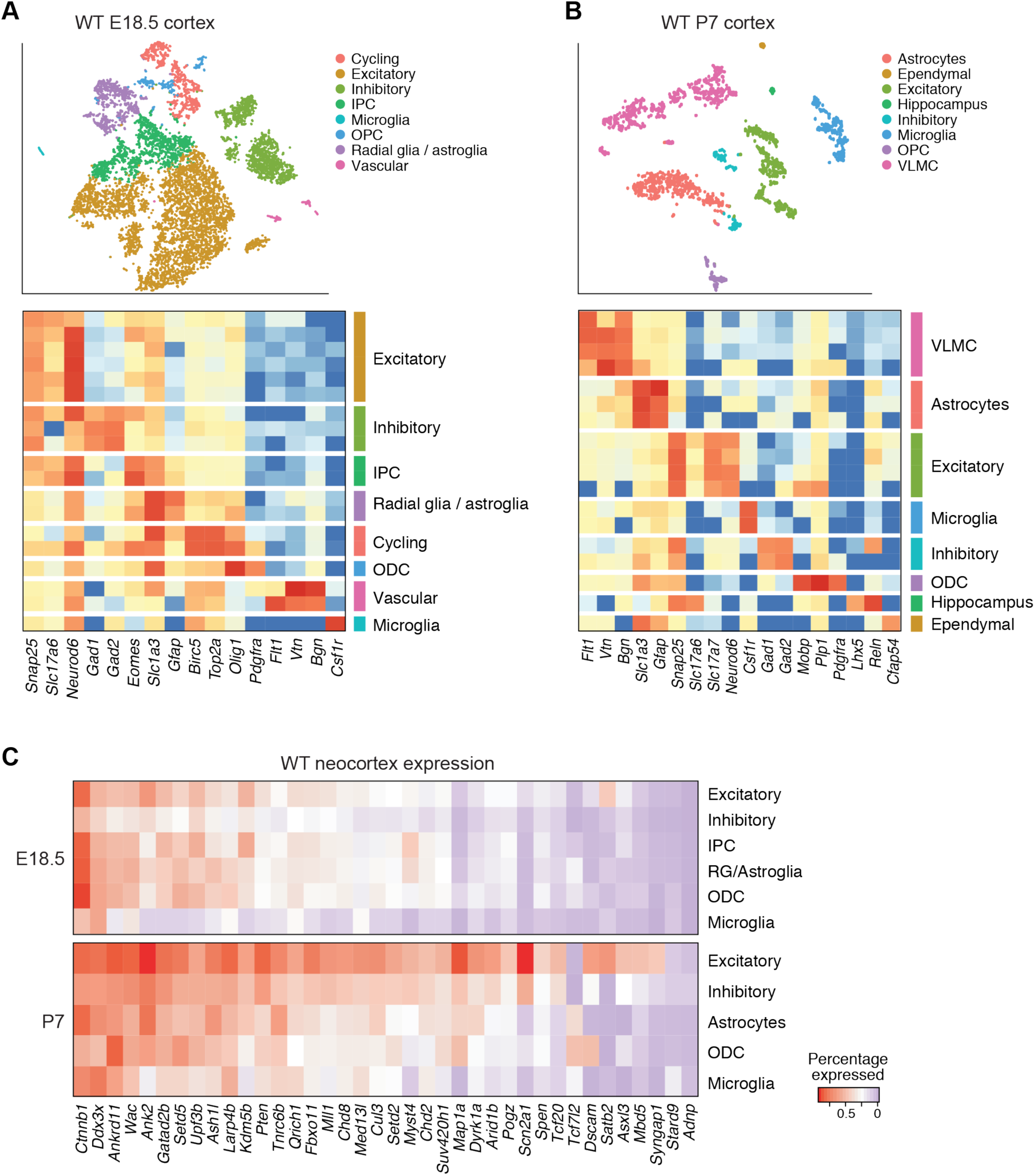
(A-B) Cell type clusters from E18.5 (public data from 10x Genomics) and WT P7 (data generated from this work) neocortex, as well as expression of cell type marker genes across identified cell clusters. (C) Expression of the 38 initially-selected risk-associated genes in the cell clusters from E18.5 and P7 wildtype cortex.

**Figure S3.**
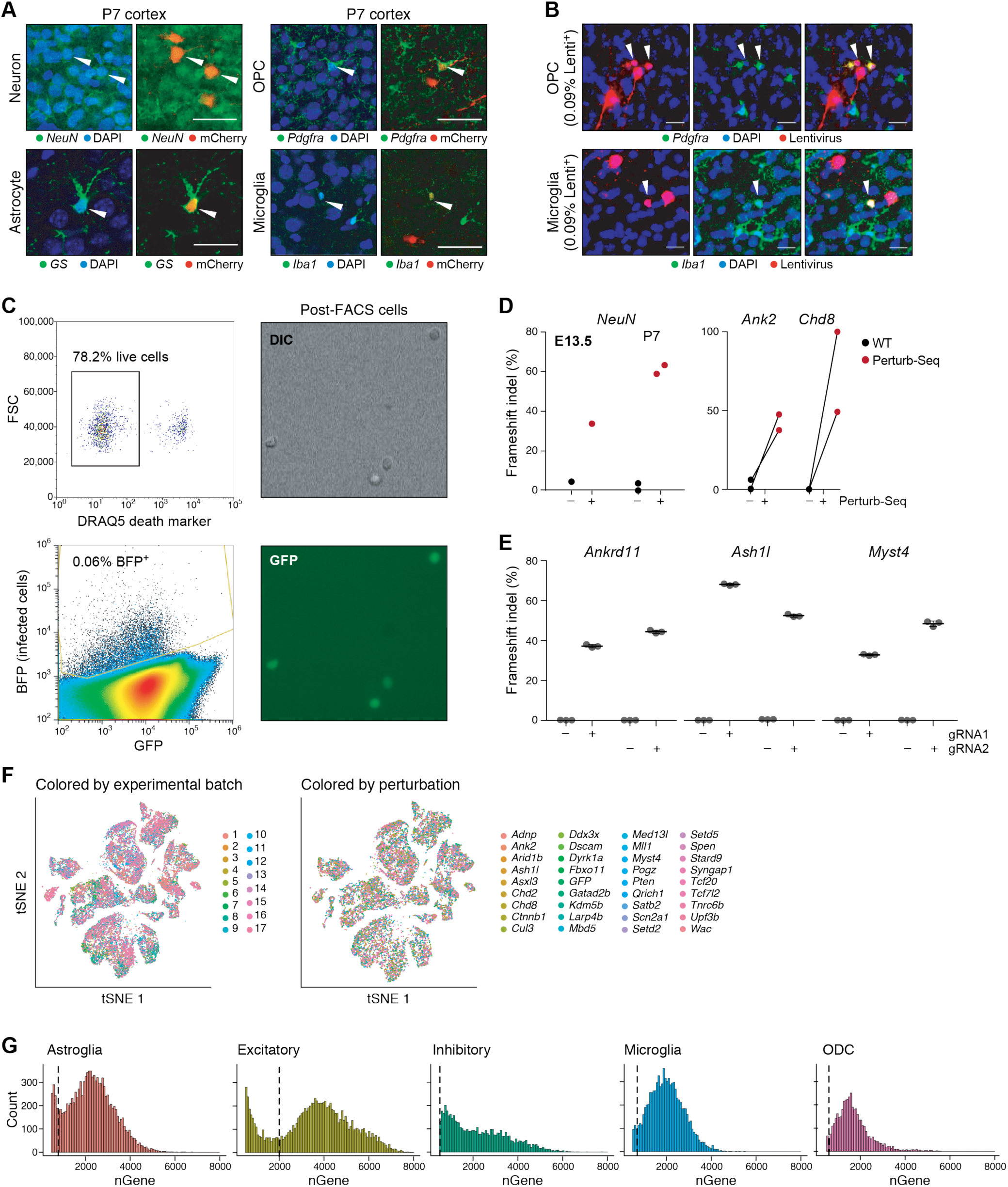
(A-B) Lentiviral injection at E12.5 can sparsely infect neurons (NeuN^+^), astrocytes (Glutamine Synthase [GS]^+^), oligodendrocyte precursor cells (PDGFRA^+^), and microglia and macrophages (IBA1^+^) in the P7 neocortex. Scale bar is 50μm. *In vivo* Perturb-Seq lentiviral vector with an mCherry expression allows immunohistochemistry and identification of the targeted cell types. (C) The proportion of live cells after FACS purification is 78.2%, and <0.1% of total dissociated cortical cells are BFP^+^. (D-E) Frameshift insertion/deletion rates of the targeted loci by CRISPR/Cas9 genome editing (D) in the infected cells *in vivo*, and (E) in mouse embryonic stem cells *in vitro*, for each gRNA. (F) Five major cell types from the Perturb-Seq cells, composed of 17 different libraries (independent experimental batches) (left) and representing 35 different perturbation groups (right). (G) Number of genes detected in each cell type in the Perturb-Seq single-cell RNA-Seq data. Quality control cutoffs for each cell type are marked by black vertical bars.

**Figure S4.**
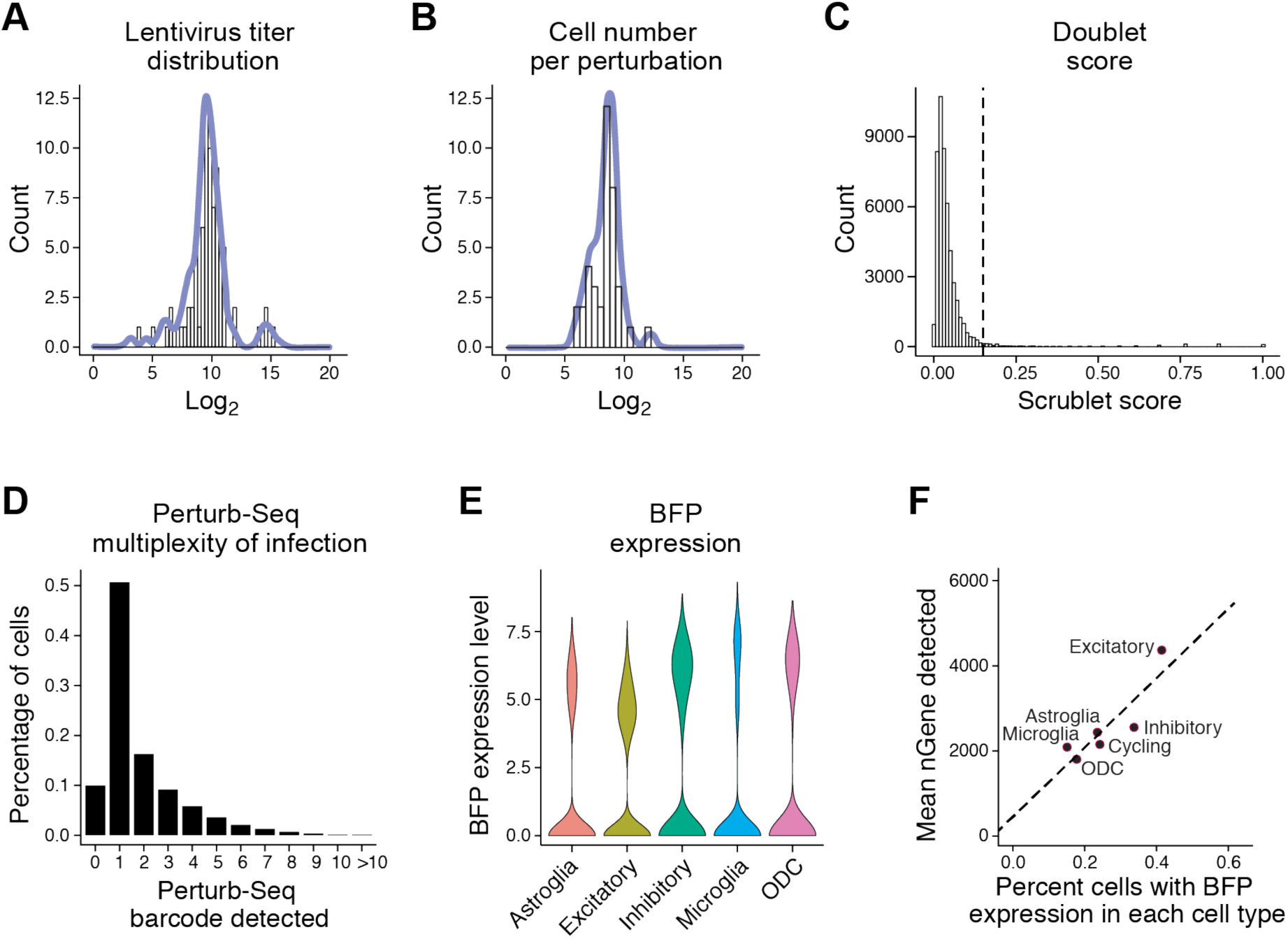
(A) The distribution of each perturbation vector in the lentiviral pool. (B) The distribution of cell numbers from each ASD perturbation group. (C) Estimated doublet score in the Perturb-Seq data using the Scrublet package; the black vertical bar represents the cutoff above which a “cell” is declared as a doublet. (D) The distribution of the number of perturbation barcodes detected per cell. (E) BFP is one of the genes with the highest expression level, detected in all five cell types. (F) BFP expression level is correlated with the number of genes detected in each cell type.

**Figure S5.**
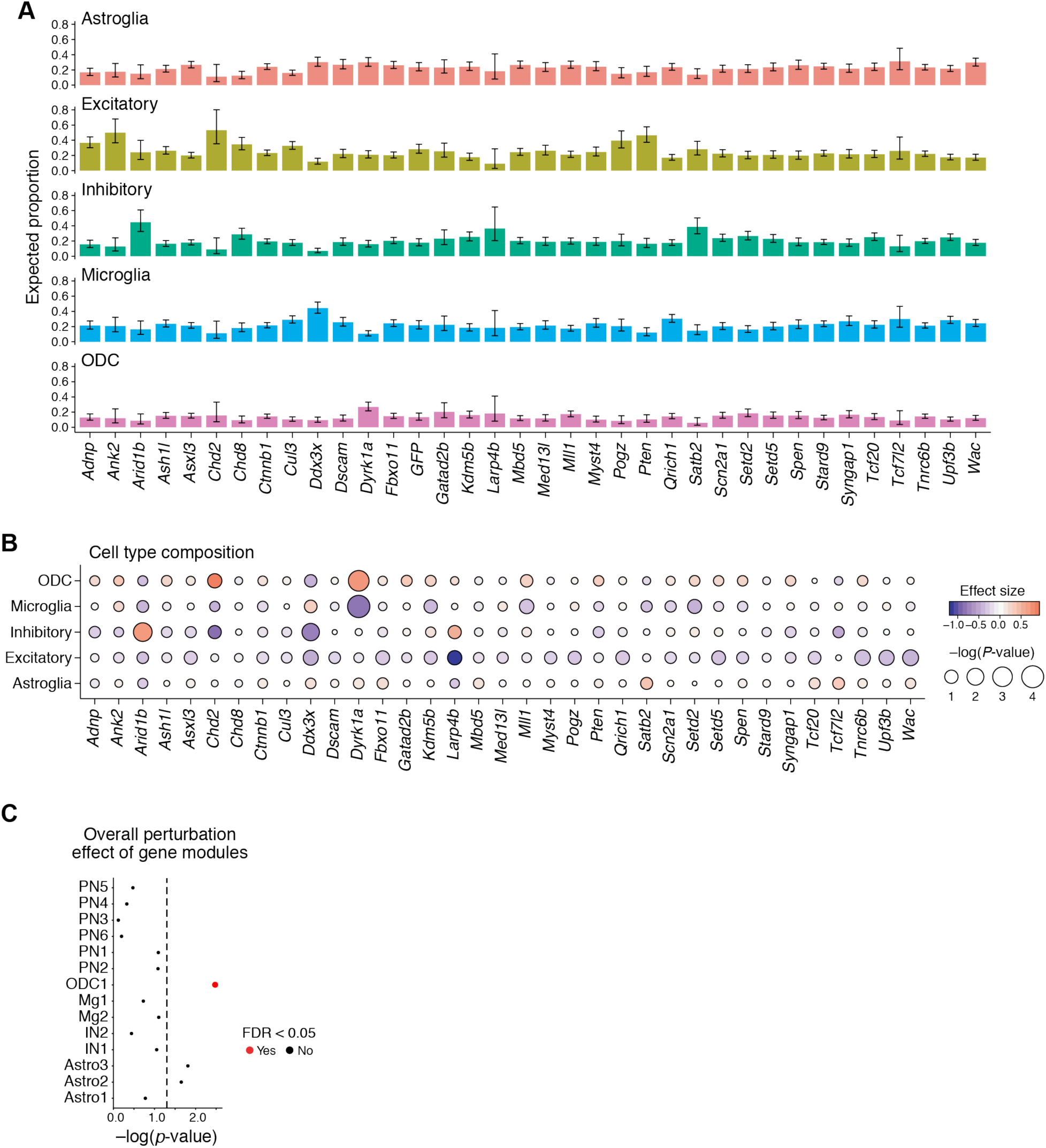
(A) Proportion of the five major cell types in each perturbation group. (B) Poisson regression for differences of cell type composition compared to the GFP control group. The size of the dots corresponds to log p-value, the color to effect size. (C) Nonparametric ANOVA analysis shows that perturbation status explains a significant portion of the variation in one module, ODC1, and nominally significant amounts in Astro2 and Astro3, all of which are identified from glial clusters.

**Figure S6.**
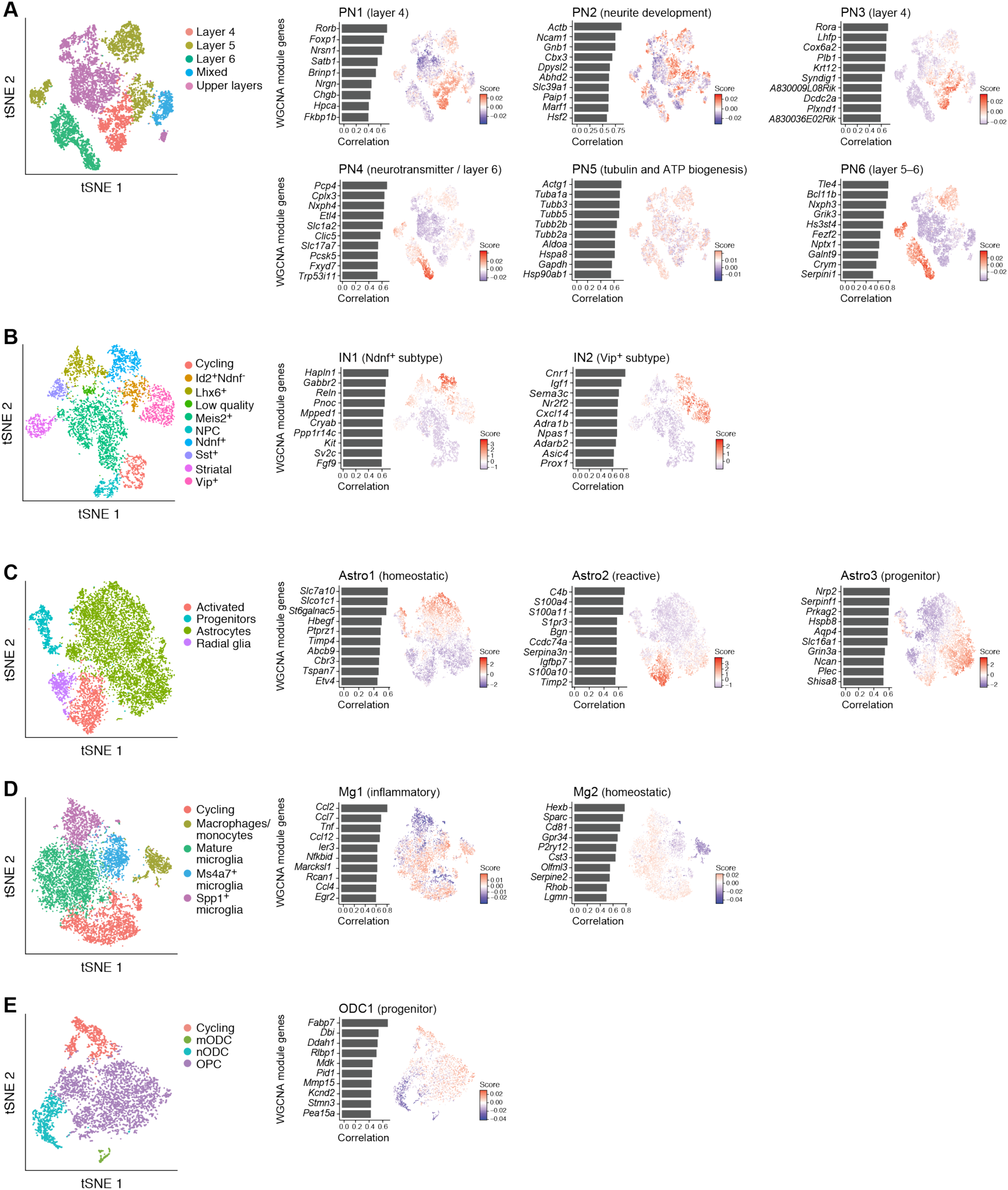
Subtypes of each major cell cluster, and feature plots of scores of gene modules identified by WGCNA, labelled by associated cell subtypes or biological processes.

**Figure S7.**
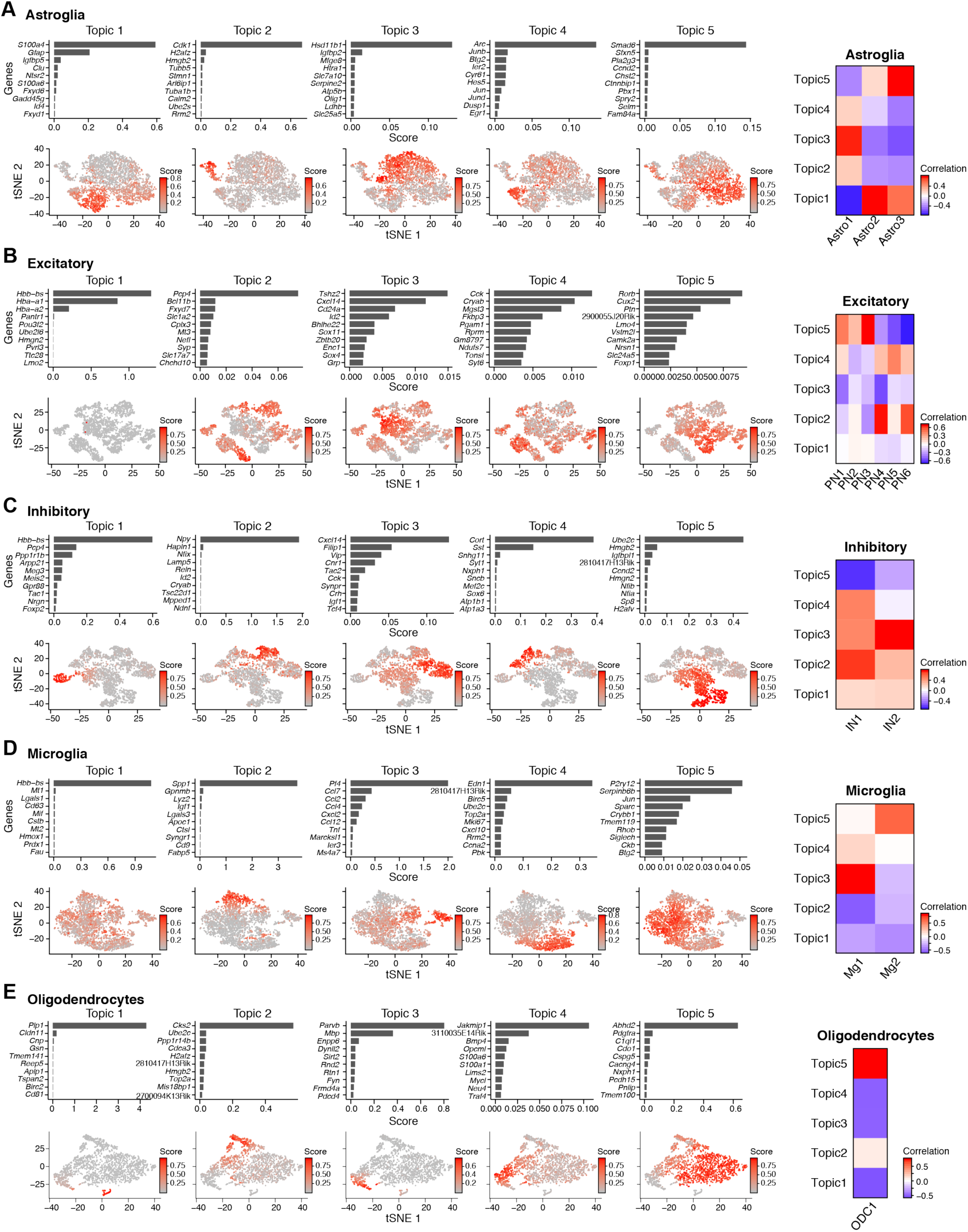
Modules identified by structural topic modeling (STM) and their correlation with WGCNA modules. Gene score indicates the lift score from STM analysis; a gene with high gene score means it is highly representative of the given topic.

**Figure S8.**
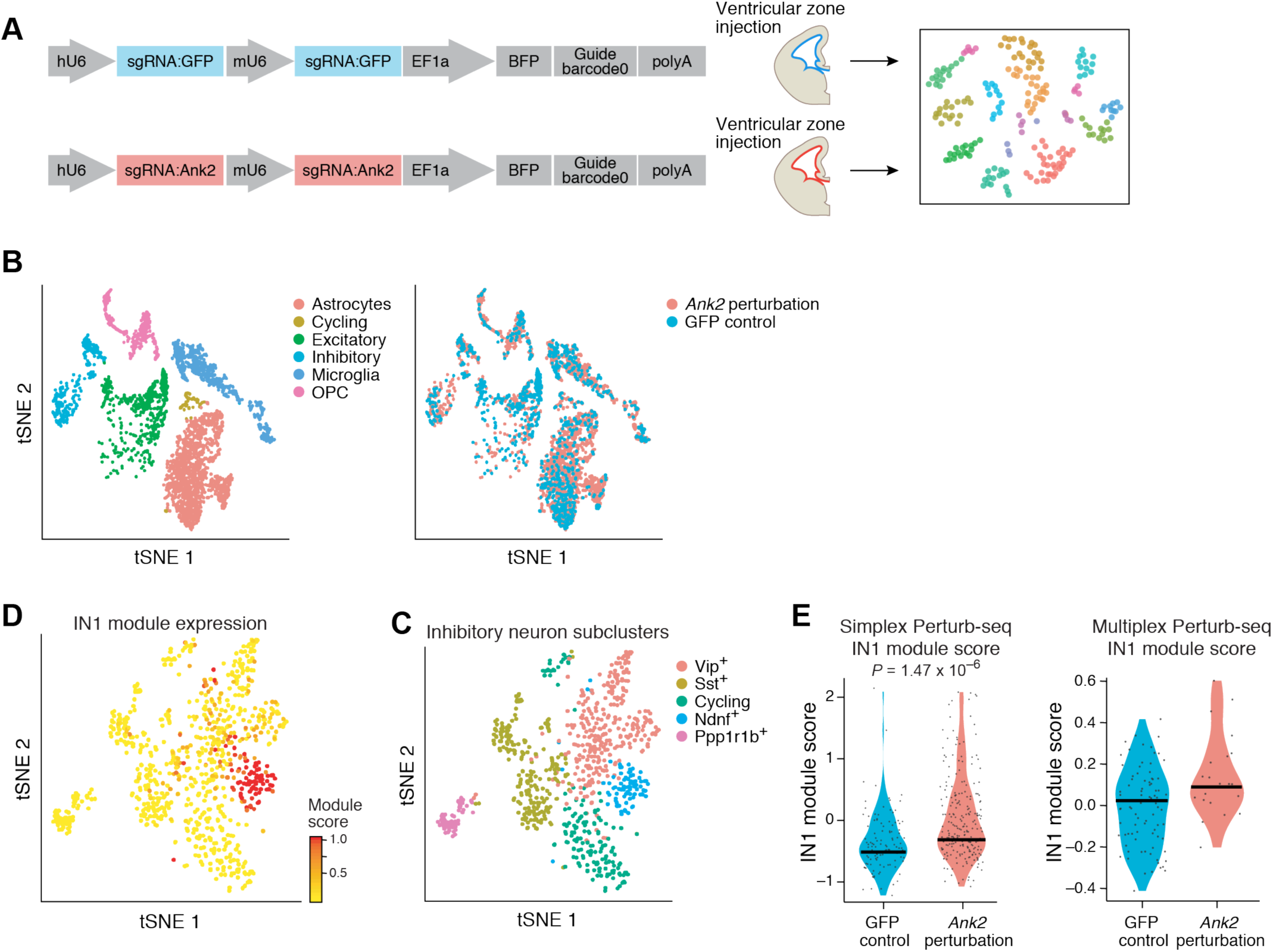
(A) Schematics of simplex Perturb-Seq of the GFP control and the ASD risk gene *Ank2*. (B) Cell type clusters from P7 neocortical simplex *Ank2* Perturb-Seq. (C) Subtype clusters of inhibitory neurons from the simplex *Ank2* Perturb-Seq. (D-E) Simplex dataset expression of the gene module IN1 identified in the pooled Perturb-Seq analysis.

**Figure S9.**
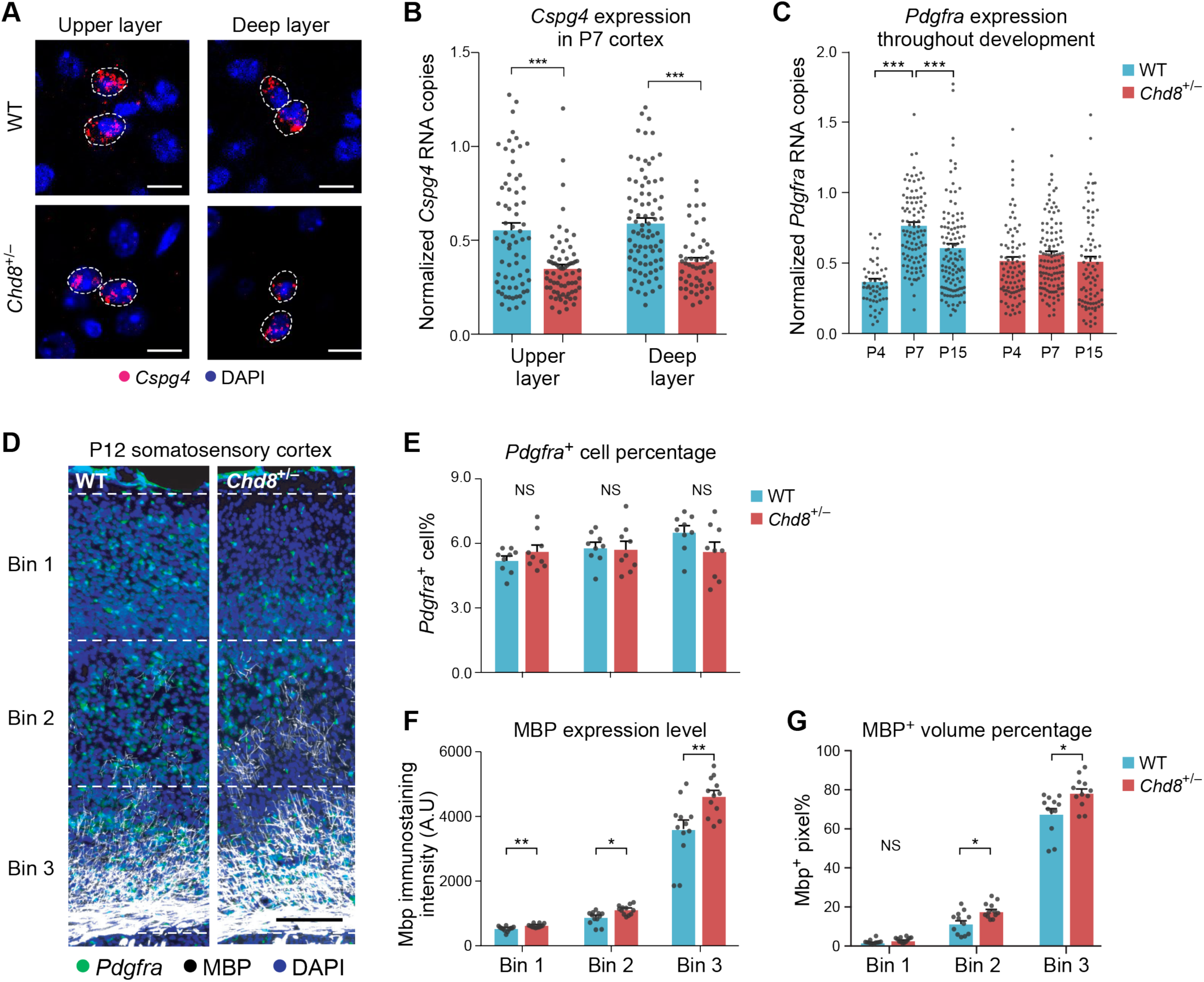
(A) *In situ* hybridization for *Cspg4*, a gene in the ODC1 gene module and a marker of oligodendrocyte precursor cells, in the somatosensory cortex of P7 *Chd8*^*+/-*^ animals and wild-type littermates. White dotted lines indicate individual *Cspg4*-positive nuclei. Scale bar is 50μm. (B-C) Quantification of *Pdgfra* expression in P4, P7, and P15 somatosensory cortex of *Chd8*^*+/-*^ and wildtype littermates. Each dot represents the gene expression measurement from one cell; error bars represent standard error of the mean. (n=2-3 animals per genotype) (D-G) Immunohistochemistry of PDGFRA and MBP, markers for immature OPC and mature oligodendrocyte, and their quantification in the somatosensory cortex of P12 *Chd8*^*+/-*^ and wild-type littermates (n=3 animals per genotype). Scale bar is 250μm.

**Figure S10.**
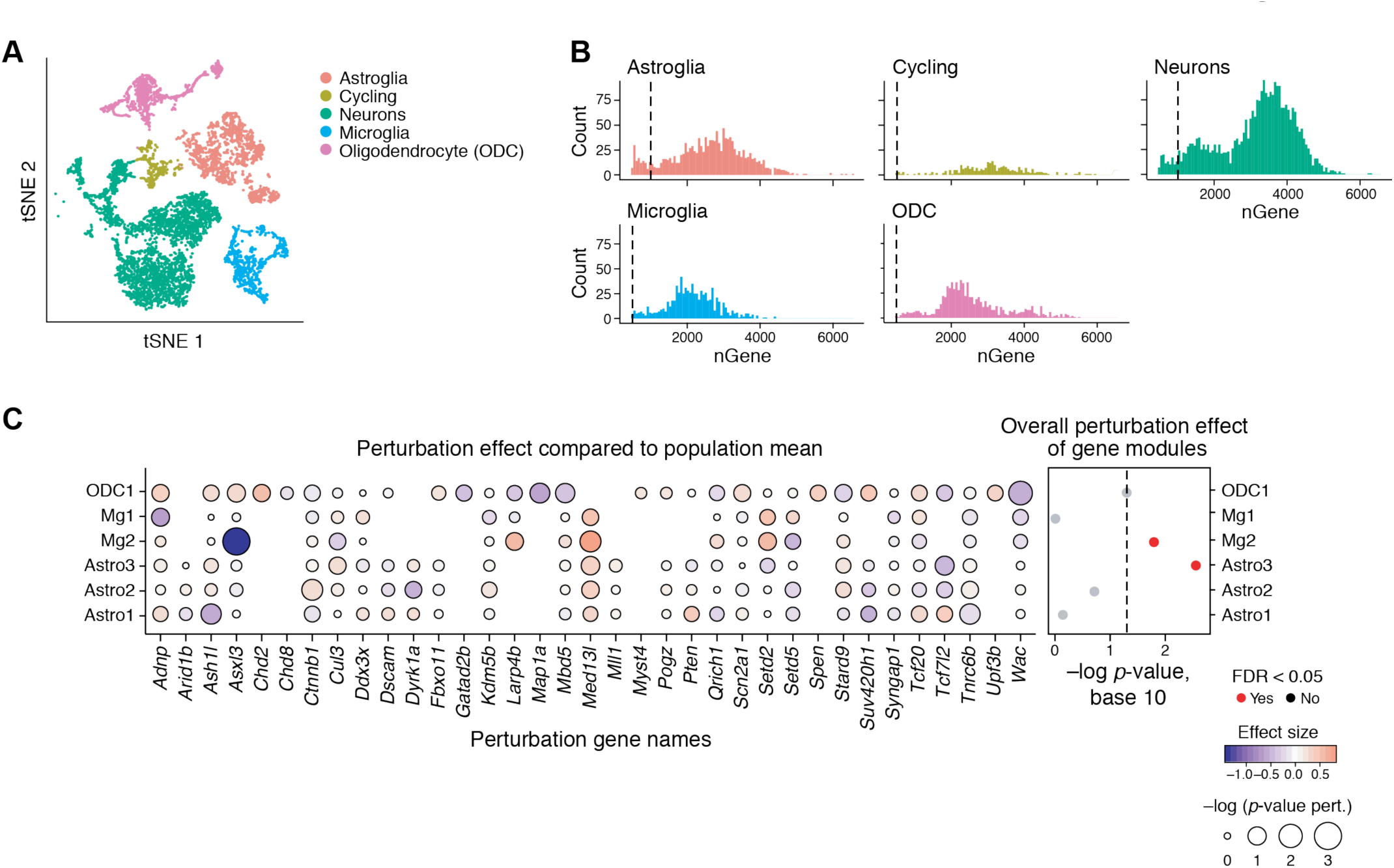
(A) Cell clusters from the striatal Perturb-Seq. (B) Number of genes detected in each cell type in the striatal Perturb-Seq data. (C) Left: Effect size for ASD risk gene perturbations in the striatum on the glial WGCNA modules identified in the cortical data, compared to the population mean. Size corresponds to log p-values, color corresponds to effect size. Right: ANOVA test identified modules significantly affected by the ASD perturbations, taken as a group, in the striatal dataset.

## Methods

### Lentiviral vector construction and production

The lentiviral vector was constructed following the Perturb-Seq publications (Adamson et al., 2016; Dixit et al., 2016; Jaitin et al., 2016). The backbone contains antiparallel cassettes of two gRNAs (Table S6) under mouse U6 and human U6 promoters, and EF1a promoter to express puromycin, BFP, and a polyadenylated barcode unique to each perturbation. Cloning of the 38 vectors were done individually. Association of each gRNA set and perturbation barcode was established by Sanger sequencing. The gRNA designs were designed using the online tool at benchling.com (Doench et al., 2016) (Table S6). Each lentivirus was packaged individually with the V2 helper plasmids (Joung et al., 2017), and the functional titer was measured individually through HEK293 cell infection and FACS measurement of BFP^+^ population before pooling equally for ultracentrifugation. The functional titer of the final lentivirus was > 5 × 10^9^ U/mL for *in utero* ventricular injection and transduction.

### *In vivo* Perturb-Seq experiment

This analysis comprises 18 independent libraries of Perturb-Seq cells. *In utero* lentiviral injection into the ventricular zone was performed at E12.5 in Cas9 transgenic mice (Platt et al., 2014), and each library was made by combining the BFP^+^ cells from 1-3 litters (4-20 animals) of P7 animals harvested on the same day.

P7 mice were anesthetized and sacrificed by decapitation, then disinfected with 70% ethanol and decapitated. The brains were quickly extracted into ice-cold PBS and cortices were micro-dissected in ice-cold Hibernate A medium (BrainBits, #HA-Lf) with B27 supplement (ThermoFisher, #17504044) under a dissecting microscope. Tissue dissociation was performed with the Papain Dissociation kit (Worthington, #LK003152). Cortices were transferred into ice-cold papain solution with DNase in a cell culture dish and cut into small pieces with a blade. The dish was then placed onto a digital rocker in a cell culture incubator for 30 mins with rocking speed at 30 rpm at 37°C. The digested tissues were collected into a 15 mL tube with 5 mL of EBSS buffer (from the Worthington kit). The mixture was triturated with a 10 mL plastic pipette 20 times and the cell suspension was carefully transferred to a 15 mL tube. 2.7 mL of EBSS, 3 mL of reconstituted Worthington inhibitor solution, and DNAse solution were added to the 15 mL tube and mixed gently. Cells were pelleted by centrifugation at 300 g for 5 mins at RT. Cells were resuspended in 0.5 mL ice-cold Hibernate A with B27 supplement (ThermoFisher, A3582801) and 10% fetal bovine serum (FBS) and subjected to FACS purification. The FACS collected cells were sorted in cold Hibernate A/B27 medium with 10% FBS (VWR, #97068). After collection the cells were centrifuged and resuspended in ice-cold PBS with 0.04% BSA (NEB, B9000S) for single-cell RNA sequencing library preparation (10x Genomics v2 chemistry). We performed the FACS purification and resuspension within 1.5 h while keeping the cells on ice to prevent necrosis, a crucial step for this experiment.

### Shared perturbation phenotypes between cortical and striatal glial cells

In order to determine whether changes in module expression observed in neocortical cells may also occur in cells of other brain regions, we tested whether the glia modules identified by *in vivo* Perturb-Seq in the neocortex were also affected in striatal glia cells. To this end, we collected striatal samples at P7, following Perturb-Seq lentiviral injection at E12.5, from 6 independent batches that passed quality control, and a total of 7,118 cells (5,933 of which are in glia clusters) with a median detection rate of 2,972 genes per cell. As before, we used Louvain clustering and known markers to identify and annotate the major cell categories as neurons, astrocytes, oligodendrocytes, and microglia (**Figure S10, Table S5**), as well as a cluster of cycling cells comprised of both interneuron progenitors and astroglia cells. Given the small number of cells in this test dataset, we included perturbations in the analysis if they had > 5 perturbed cells per perturbation group, recognizing that this could increase the noise in effect estimates.

Using the same nonparametric ANOVA approach, we find that perturbation status explained a significant amount of the total variation in the astrocyte progenitor module (Astro3) in the striatal dataset, similar to the cortical dataset (FDR corrected *P*<0.01) (**Figure S10, Figure 2F**). The ODC1 module is nearly significant (*P*<0.06, FDR corrected *P*<0.11), likely reflecting limited power due to the small number of ODC cells profiled. Moreover, the direction of effect of most individual gene perturbations largely agreed with those in the cortical data. In particular, both *Gatad2b* and *Chd8* perturbations showed decreased expression of the ODC1 module in this striatal data, as observed in the cortex (**Figure S10**). Thus, at least some of the ASD gene perturbation effects appear to generalize in cells of more than one brain region.

### RNA *in situ* hybridization

RNAscope fluorescent *in situ* hybridization was performed on fixed-frozen tissue. Mice were anesthetized and transcardially perfused with ice-cold PBS followed by ice-cold 4% paraformaldehyde in PBS. Dissected brains were postfixed overnight in 4% paraformaldehyde at 4°C, and cryoprotected in 30% sucrose. Brains were then embedded in optimal cutting temperature (OCT) compound (Tissue-Tek, #4583) and 15-20μm tissue sections were prepared. Multiplex RNAscope v1 was performed based on manufacturer’s instructions. Probes against the following mRNA were used: *Pdgfra, Cspg4*, and *Fezf2* (ACDBio). Quantification were performed by StarSearch; gene expression copy number were normalized to pixel area (https://www.seas.upenn.edu/~rajlab/StarSearch/launch.html).

### Immunohistochemistry

Mice were anesthetized and transcardially perfused with ice-cold PBS followed by ice-cold 4% paraformaldehyde in PBS. Dissected brains were postfixed overnight in 4% paraformaldehyde at 4 °C, and cryoprotected in 30% sucrose. The brains were embedded in OCT compound (Tissue-Tek, #4583) and 15-20μm tissue sections were prepared. The slides with tissue sections were incubated with blocking media (6% donkey serum in 0.3% Triton with PBS) for 1hr, then incubated with primary antibodies in a 1:3 dilution of blocking media in PBS with 0.3% Triton overnight at 4 °C. Slides were washed with PBS with 0.3% Triton 4 times to remove the excess primary antibody. Secondary antibodies were applied at 1:800 dilution in blocking media and incubated for 2hr at room temperature. Slides were then washed 4 times with PBS with 0.3% Triton, and incubated with DAPI for 10 mins before mounting with Fluoromount G (Invitrogen, #00-4958-02). The antibodies and dilutions were: Mouse anti-NeuN antibody (mab377, 1:500; Millipore), Mouse anti-GS antibody (mab302, 1:500; Millipore), Goat anti-Pdgfra antibody (AF1062, 1:200; R&D System), Rabbit Iba1 antibody (019-19741, 1:400; Wako), Chicken anti-GFP antibody (ab16901, 1:500; Millipore), Mouse anti-Satb2 (ab51502, 1:50; Abcam), Rat anti-Ctip2 (ab18465, 1:100, Abcam), Rabbit anti-Sox6 (ab30455, 1:500; Abcam), Rat anti-Mbp (mab386, 1:100; Millipore). All images were acquired using either a custom-built spinning disk confocal microscope equipped with image acquisition NIS-Elements software, or a Carl Zeiss epifluorescent microscope with Zen software.

### Perturb-Seq profiling

Single-cell RNA sequencing libraries were created using the Chromium Single Cell 3’ Solution v2 kit (10x Genomics) following the manufacturer’s protocol. Each library was sequenced with Illumina NextSeq high-output 75-cycle kit with sequencing saturation above 70%. Reads were aligned to the mm10 mouse genome reference using the Cell Ranger package (10x Genomics).

To sequence the perturbation barcode, dial-out PCR was performed to extract the perturbation barcode in each cell. This is modified from Dixit et al (Dixit et al., 2016) to be compatible with the 10x Genomic V2 chemistry instead of V1. The PCR product was sequenced with the 10x libraries, and demultiplexed to extract the perturbation information.

#### Forward primer

CAAGCAGAAGACGGCATACGAGAT-TCGCCTTA-GTCTCGTGGGCTCGGAGATGTGTATAAGAGACAG-TAGCAAACTGGGGCACAAGC

#### Reverse primer (i5)

AATGATACGGCGACCACCGAGATCTACAC

## Data Analysis

### Data pre-processing

BCL files were transformed into fastq files using the cellranger mkfastq command, using CellRanger V2.1.0. Bam files and expression matrices were generated from these fastq files using the cellranger count command, using force_cells=8000.

### Identification of perturbation barcode

In order to extract perturbation information from the dial-out reads, we modified code from the original Perturb-Seq paper (Dixit et al., 2016) to work with 10x V2 chemistry, and applied it to our data (original code at https://github.com/asncd/MIMOSCA). This resulted in a cell-by-perturbation UMI count matrix. To extract perturbation information from the 10x reads, a fasta file was first created with one entry for each perturbation, containing the sequence of the perturbation barcode and the surrounding sequence. This fasta file was turned into a STAR reference (Dobin et al., 2013), referred to as the PBC reference. Unmapped reads containing either AGAATT or CCTAGA as a subsequence were extracted from the Cell Ranger bam file, and then mapped to this new reference. Low quality reads were filtered out using the following filters: (i) used “samtools view - F 2820” to filter out unmapped, multimapped, and low quality reads from the PBC mapped bam file, (ii) removed reads with quality scores <255, (iii) removed reads whose 5’ end did not map between 655 and 714bp into the PBC reference, to help exclude reads that did not overlap enough bases in the perturbation barcode for proper identification of the perturbation, and (iv) removed reads whose edit distance from the PBC reference was >2. Reads were then assigned to the perturbation they mapped best. Cell barcodes and UMIs were extracted, and a cell-by-perturbation UMI count matrix was created. This matrix was used to assign cells to perturbations in the same way as with the dial-out data. As with the dialout data, if a cell had one perturbation with >1.3x the number of UMIs assigned to it than the next best perturbation based on the 10x sequence, that cell was assigned to that perturbation in the 10x data; otherwise, the cell was declared to have multiple perturbations. We then only kept cells for which either i) the assigned 10x and dialout perturbations agree or ii) the cell was assigned to a perturbation by one method but not assigned to a perturbation in the other.

### Cell type clustering analysis

UMI count data was loaded into R and processed using the Seurat v 2.2 package (Butler et al., 2018). Data were scaled to counts per million and log normalized. Cells expressing less than 500 genes were removed. Variable genes were found using FindVariableGenes with x.low.cutoff=1 for each batch separately. Genes that were found to be variable in at least 4 batches were combined into a final combined list of variable genes. The normalized data was scaled with ScaleData on the variable genes, regressing out the effects of nUMI, and PCA was performed. Clustering was performed with the FindClusters function (with default parameters, except for resolution=1.2 and using 28 PCs). tSNE plots were generated with RunTSNE (RunTSNE (with default parameters, except with 28 PCs and pca=F). Clusters were assigned to cell types based on marker genes from the literature, mousebrain.org (Zeisel et al., 2018), and DropViz (Saunders et al., 2018). For each cell type a more refined nGene cutoff was identified (**Figure S3**), and cells of that cell type with less than that filter were removed from further consideration. We focused only on cells of 5 key types (projection neurons, inhibitory neurons, oligodendrocytes, microglia, and astroglia) and removed the rest.

For subclustering individual cell types, the cells of that cell type were extracted from the larger Seurat object. Variable genes were chosen as above, and data was scaled with ScaleData, regressing out the effects of nUMI and batch, followed by PCA. Clustering was performed with FindClusters (with default parameters except for varying resolutions and number of PCs, **Table S7**). tSNE was performed with RunTSNE (with default parameters, except with different numbers of PCs and pca=F).

### Testing WGCNA Gene Sets

WGCNA was performed for each cell cluster based on the published pipeline (Langfelder and Horvath, 2008). We manually removed modules that were driven by outlier cells. For a given cell type, each WGCNA gene set was input into moduleEigengenes to calculate a gene-set score for that set of genes. All cells without an assigned perturbation were removed.

Linear regression was used to test the relationship between perturbations and WGCNA gene scores, correcting for batch and number of genes with the lm function in R, using the formula:

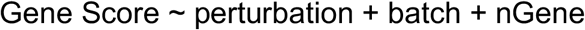

Associated p-values and effect sizes were extracted. In addition, a permutation-based approach was used to calculate an empirical p-value to ensure the model-based p-values reported by lm were accurate. Specifically, the perturbation labels of cells were randomly permuted within each batch, and the absolute effect size for each perturbation was calculated as above on this permuted data. This was repeated 10,000 times. The empirical p-value was the proportion of permutations (including the original data) with absolute effect size larger than that of the original data. FDR correction was performed using the Benjamini & Hochberg procedure.

### Structural Topic Modelling

Structural topic modelling (STM) was performed separately on each cell type of interest using the STM package in R (Roberts et al.). Count data from cells of a given type were extracted from the Seurat object, along with corresponding meta data. Genes that occurred in <5% or >90% of cells were removed, as were mitochondrial and ribosomal genes. In addition, only genes that were expressed in at least one cell in all batches were retained in order to help reduce batch effects. The resulting count matrix was provided as input to the STM function, along with the meta data and with parameters LDAbeta=T, interactions=F. The formula used by the STM function was

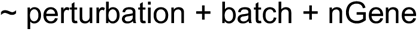

This specifies a model that assumed topic proportions were dependent on perturbation, number of genes, and batch. We ran this model on each dataset with 5 topics. Top 10 genes for each topic were extracted with the labelTopics function.

To test for correlations between perturbations and topics, the theta matrix (the matrix containing proportions of topics per cell) was extracted from the STM matrix. For each topic, linear regression was used to test how the per-cell proportions for each topic related to perturbations (after setting GFP to be the reference perturbation), correcting for nGene and batch. In particular, the lm function in R was used, with the formula:

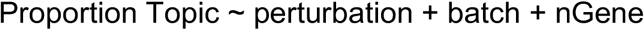

Effect sizes were extracted from the resulting lm object. An empirical p-value was calculated, as for WGCNA. FDR correction was performed using the Benjamini & Hochberg procedure.

### Cell Type Gene Expression

Expression data for the E18.5 mouse brain (9k dataset) was downloaded from the 10X website(https://support.10xgenomics.com/single-cell-gene-expression/datasets/2.1.0/neuron_9k). The WT P7 data were generated from this paper. The P7 fastq files were run through the standard Cellranger pipeline. The data from both datasets were loaded into Seurat separately and transformed to log counts per million. Cells with <500 genes were removed in both datasets. Variable genes were found using FindVariableGenes with x.low.cutoff=1, and the data was scaled with ScaleData, correcting for nUMI. PCA was performed, followed by TSNE and clustering with FindClusters. Cell types were identified with marker genes, and contaminating/vascular cell types were removed.

In each dataset MAST (Finak et al., 2015) was used to find the differentially expressed genes in each cluster, relative to all cells outside that cluster. This was done correcting for the scaled nUMI and removing genes that occurred in less than 10 cells. Average expression was calculated for each gene in each cluster.

